# Reorientation of the primary body axis by ectopic embryonic cWnt signaling

**DOI:** 10.1101/220988

**Authors:** Naveen Wijesena, Mark Q. Martindale

**Author notes:** Corresponding author: Mark Q. Martindale, Whitney Laboratory for Marine Bioscience, University of Florida, Saint Augustine, FL 32080.

## Abstract

Gastrulation is a crucial time during embryogenesis when cells make important decisions on what larval or adult tissues (i.e. ectodermal, mesodermal, or endodermal) are going to generate. The evolution of gastrulation was a pivotal event during metazoan evolution, as it paved the way for diversification of the metazoan clade from a hollow, ciliated, radially symmetrical ancestor (1, 2). The position of the site of gastrulation (that segregates internal endomesodermal precursors from outer ectodermal tissue) has played a role in our understanding patterns of body plan evolution (e.g. deuterostomes vs protostomes) and is tightly regulated during development. In bilaterians (a large clade of bilaterally symmetrical animals that represent over 99% of all extant species), the site of gastrulation is determined by a localized molecular asymmetry resulting from a differential distribution of maternal determinants (3) along the so-called animal-vegetal axis where the animal pole is marked by the site of polar body release during meiosis (2, 4).

In most bilaterians, the site of gastrulation (endomesoderm formation) is generated from blastomeres derived from the vegetal pole (5), however, in cnidarians (e.g. corals, sea anemones, and “jellyfish) (6, 7), the sister group to all bilaterians and ctenophores (e.g. comb jellies), likely to be the earliest branching group of extant metazoans (8), gastrulation occurs at the animal pole (6, 9). These observations suggest that gastrulation and endoderm formation originally evolved at the animal pole leading to the formation of a gut with a single opening (oral) in the metazoan ancestor and that gastrulation later shifted to the vegetal pole in the last common ancestor of the bilaterian lineage. Molecular evidence for such a switch in the site of gastrulation comes from the fact that the site of gastrulation in both bilaterians (10) and non bilaterians (11) is marked by the site of nuclearization of the protein βcatenin, which is a downstream target of the canonical Wnt/ βcatenin (cWnt) signaling pathway. The causal role of localized activation of cWnt signaling in determining axial properties of the embryo and the adult suggests that a change in the site of localized activation of cWnt signaling from non-bilaterians to bilaterians could have resulted in a change in the site of gastrulation and endomesoderm specification. We tested this hypothesis by mis-activating cWnt signaling in the developing embryos of the anthozoan cnidarian, *Nematostella vectensis*. We show that ectopic activation of cWnt signaling at a site other than the animal pole results in a new oral-aboral axis specified by the new site of cWnt activation.

We used a two-step experimental approach to mis-activate cWnt signaling in the developing *N. vectensis* embryos (Figure 1). First, we “erased” the endogenous oral-aboral axis by inhibiting the activation of cWnt signaling in the embryo by injecting zygotes with a mRNA coding for a dominant-negative form of the Dishevelled protein, DshDIX::GFP to (12). Then at the 16-cell stage, a random blastomere was injected with a mixture of two mRNAs, an “activated” form of βcatenin that is immune to targeted degradation (11) and the complete ORF of the βcatenin binding factor Tcf (Tcf::venus) to ectopically activate cWnt signaling at a site other than the animal pole. Injection of only the activated form of βcatenin did not result in a new axis (Supplementary Figure 1) and this could be due to selective localization of Tcf to the animal pole blastomeres of developing *N. vectensis* embryos (13). Compared to control embryos at 30 hours post fertilization, injected embryos had undergone normal primary archenteron invagination but showed defects in cell fate specification (Figure 1). Because cell shape changes associated with primary archenteron invagination is regulated by Wnt/PCP signaling in *N. vectensis* (3), the site of primary archenteron invagination remains unchanged in experimental embryos and thus marks the original position of the animal pole (Figure 1). In control embryos, the oral marker *NvBrachyury* is expressed at the blastopore and the aboral marker *NvSix3* is expressed in the aboral domain opposite the blastopore (Figure 1). In contrast, experimental embryos with ectopic activation of cWnt signaling showed ectopic expression of *NvSix3* in regions opposite to the injection site and *NvBrachyury*, a downstream target of βcatenin/Tcf signaling (14) expression shifted from the blastopore to a site opposite *NvSix3* expression (Figure 1).

**Figure 1:**
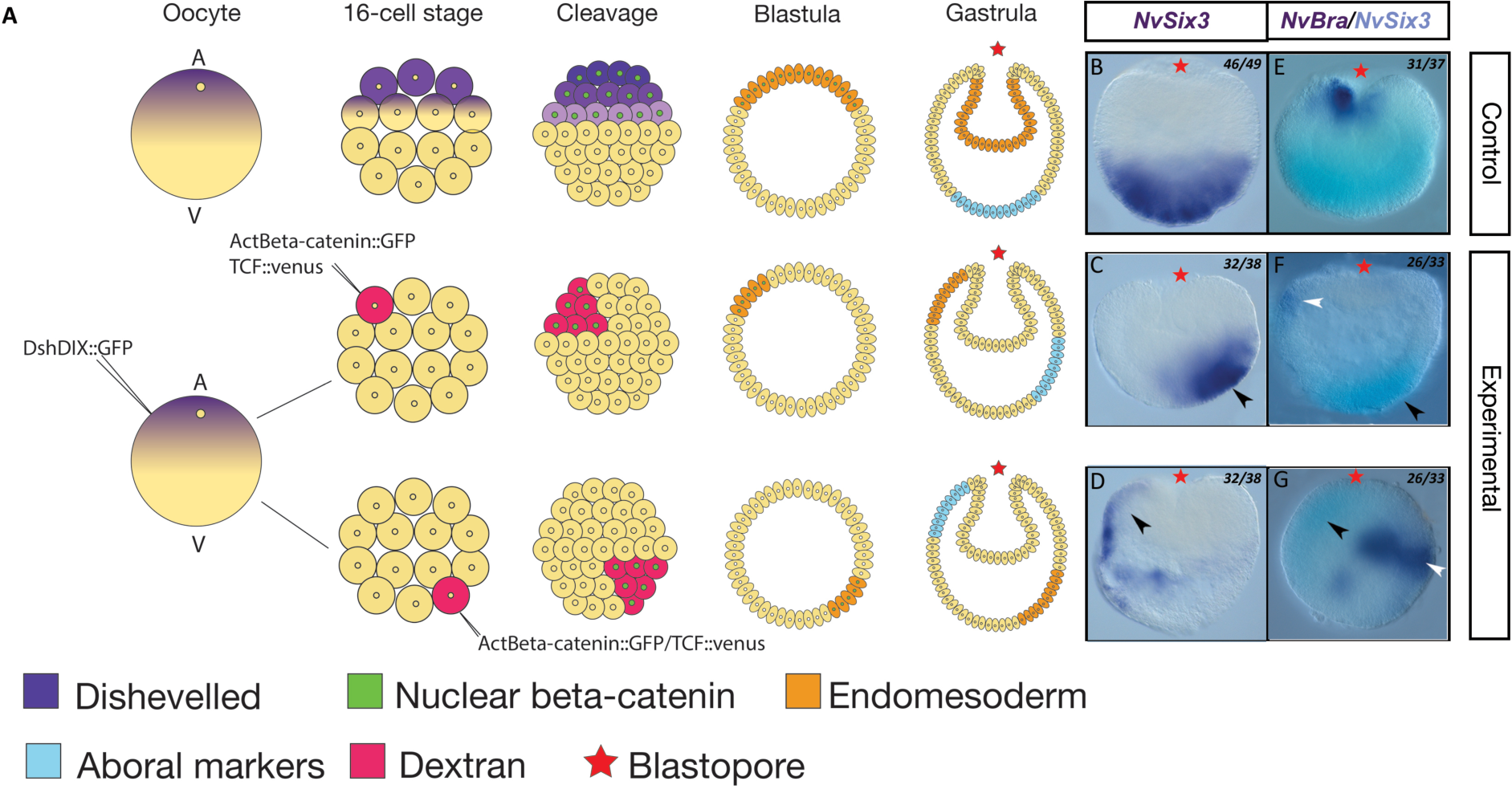
Ectopic activation of cWnt signaling in N. vectensis embryos results in a new oral-aboral axis. Diagram showing the experimental design where cWnt signaling is activated in an ectopic site after inactivation of endogenous cWnt signaling (A). WMISH of *NvSix3* shows the ectopic expression of *NvSix3* (black arrow heads) (C, D) in embryos where cWnt signaling is activated in an ectopic site compared to control embryos (B). WMISH of *NvSix3* and *NvBra* shows the ectopic expression of *NvBra* (white arrow heads) at a site other than the blastopore and *NvSix3* (black arrow heads) is expressed opposite the site of *NvBra* expression (F,G) compared to control embryos (E).

Our data show that this change in the site of gastrulation and endomesoderm specification is the result of moving localized components of the Wnt signaling pathways to the vegetal pole in the last common ancestor of all bilaterians (Supplementary Figure 2). It has been shown experimentally in both cnidarians and ctenophores that moving the zygotic nucleus from the animal pole to an ectopic site completely re-specifies the oral-aboral axis (6, 15), and in two cnidarians the βcatenin stabilizing protein Dsh is associated with the female pronucleus prior to first cleavage (12). This shows that, unlike most bilaterians, in which the definitive embryonic (animal-vegetal) and organismal axial properties are stablley established maternally, in ctenophores and cnidarians, the relationship between the AV axis and the axial properties of the embryo only establish the position of the selective activation of Wnt/PCP and cWnt signaling through localized components of the Wnt signaling pathways.

These data provide a plausible mechanism for the change in the site of gastrulation from the animal pole in cnidarians (and ctenophores?) to the vegetal pole in bilaterians (Supplementary Figure 2). The selective activation of cWnt and the module of endomesodermal gene expression that is downstream of βcatenin/Tcf (5, 14) generates endomesodermal fates at the vegetal pole. This change in the site of gastrulation released, arguably the largest developmental constraint in metazoan evolution by spatially separating endomesodermal fates to the vegetal pole from oral/neural fates derived from derivatives of the animal pole and paved the way for the diversification (e.g. cephalization) of the bilaterian nervous system.

---

**Supplementary Figure 1:**
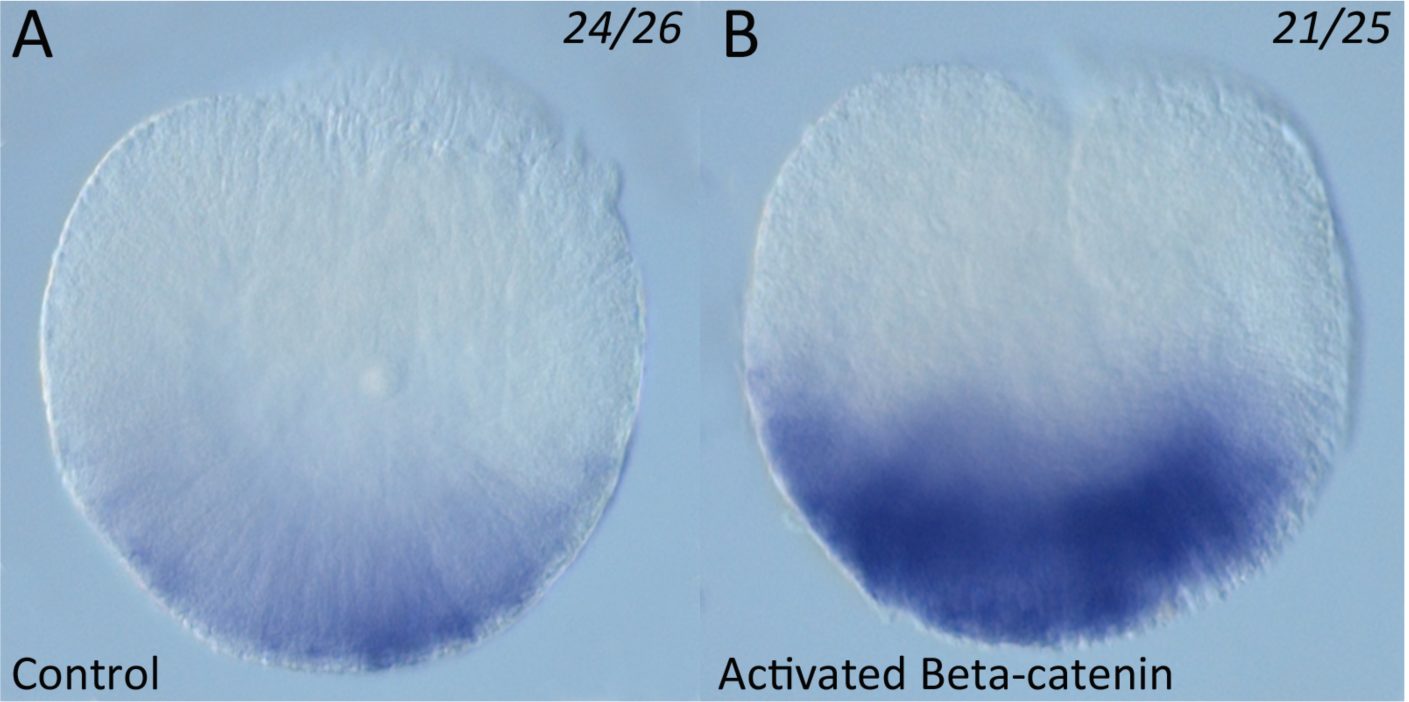
Activated Beta-catenin is not sufficient for specifying a new oral-aboral axis. Injection of activated beta-catenin by itself does not change the expression patter of *NvSix3* (B) compared to control embryos (A).

**Supplementary Figure 2:**
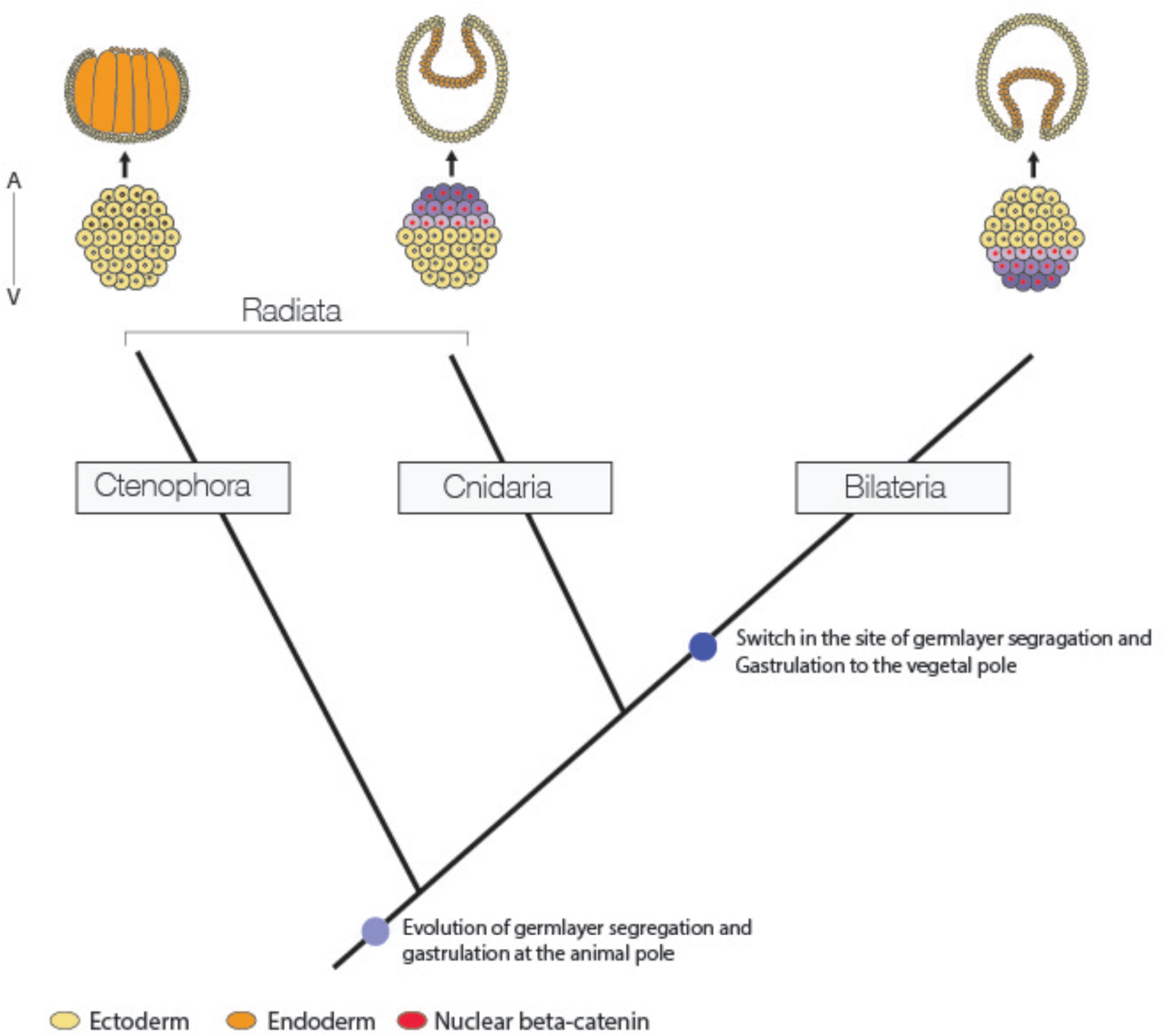
A model for the evolution of germ layer segregation and gastrulation. A molecular asymmetry present in the unicellular last common ancestor of metazoans was co-opted as a scaffold to localize maternal Wnt pathway components regulating germ layer segregation and gastrulation to the animal pole in non-bilaterians. This scaffold was moved to the vegetal pole in the in the urbilaterian, triggering extensive body plan radiation in the bilaterian clade.

